# *C. elegans* TFIIH subunit GTF-2H5/TTDA is a non-essential transcription factor indispensable for DNA repair

**DOI:** 10.1101/2021.06.04.447037

**Authors:** Karen L. Thijssen, Melanie van der Woude, Carlota Davó-Martínez, Mariangela Sabatella, Wim Vermeulen, Hannes Lans

## Abstract

The 10-subunit TFIIH complex is vital to both transcription initiation and nucleotide excision repair. Hereditary mutations in its smallest subunit, TTDA/GTF2H5, cause a photosensitive form of the rare developmental brittle hair disorder trichothiodystrophy (TTD). Some TTD features are thought to be caused by subtle transcription or gene expression defects. Strikingly, TTDA/GTF2H5 knockout mice are not viable, which makes it difficult to investigate how TTDA/GTF2H5 promotes transcription *in vivo*. Here, we show that deficiency of the *C. elegans* TTDA ortholog GTF-2H5 is, however, compatible with viability and growth, in contrast to depletion of other TFIIH subunits. We also show that GTF-2H5 promotes the stability of TFIIH in multiple tissues and is indispensable for nucleotide excision repair, in which it facilitates recruitment of the TFIIH complex to DNA damage. Strikingly, when transcription is challenged, *gtf-2H5* embryos die due to the intrinsic TFIIH fragility in the absence of GTF-2H5. These results support the idea that TTDA/GTF2H5 mutations cause transcription impairment underlying trichothiodystrophy and establish *C. elegans* as potential model for studying the pathogenesis of this disease.

## Introduction

The general transcription factor TFIIH is an evolutionary conserved 10-subunit complex that has essential functions in transcription initiation and nucleotide excision repair (NER) (Compe & Egly, 2016). It consists of a core complex comprised of subunits XPB/ERCC3, XPD/ERCC2, p62/GTF2H1, p44/GTF2H2, p34/GTF2H3, p52/GTF2H4 and p8/TTDA/GTF2H5 and the cyclin-activating kinase (CAK) complex consisting of MAT1/MNAT1, CDK7 and Cyclin H/CCNH (Luo *et al*, 2015). In RNA polymerase II (Pol II)-driven transcription initiation, TFIIH binds the promoter and, stimulated by TFIIE (Compe *et al*, 2019; Ohkuma & Roeder, 1994), facilitates promoter escape and RNA synthesis by DNA helix opening using its XPB helicase/translocase subunit (Coin *et al*, 1999; Tirode *et al*, 1999; Fishburn *et al*, 2015) and by phosphorylating the C-terminal domain of RPB1, the largest Pol II subunit, via its CDK7 subunit (Lu *et al*, 1992). TFIIH is furthermore thought to function in RNA Pol I transcription (Hoogstraten *et al*, 2002; Iben *et al*, 2002) and to regulate the recruitment and/or activity of various transcriptional regulators (Keriel *et al*, 2002; Singh *et al*, 2015). In NER, TFIIH functions in both the global genome (GG-NER) and the transcription-coupled (TC-NER) subpathway. GG-NER repairs helix-distorting single-stranded DNA lesions anywhere in the genome, whereas TC-NER repairs any lesion that blocks elongating Pol II (Marteijn *et al*, 2014; Schärer, 2013). TFIIH is recruited to lesions upon DNA damage detection and, together with NER factor XPA, opens the DNA using its XPD helicase subunit, thereby verifying the presence of damage and providing a substrate for downstream endonucleases ERCC1/XPF and XPG to cut out the damaged DNA (Sugasawa *et al*, 2009; Winkler *et al*, 2000). These essential functions of TFIIH are conserved from yeast to humans.

Mutations in the individual subunits of TFIIH cause several diseases characterized by growth and neurodevelopmental failure (Lehmann, 2003; Ferri *et al*, 2020). Mild mutations in XPB and XPD, which only affect TFIIH function in GG-NER, cause xeroderma pigmentosum, which is characterized by sun sensitivity and cancer susceptibility. More disruptive mutations in these subunits additionally cause Cockayne syndrome (CS) features, such as growth failure and progressive neurodegeneration, of which the exact pathogenesis is still debated and not fully understood. We proposed that specific impairment of TC-NER, in combination with prolonged lesion stalling of Pol II or the NER core complex, causes or contributes to CS features (Lans *et al*, 2019; Sabatella *et al*, 2018). However, other causes such as specific transcriptional or mitochondrial defects have been put forward (Wang *et al*, 2014; Karikkineth *et al*, 2017). A third group of mutations in XPB and XPD, likely those affecting also TFIIH transcription function, cause a photosensitive form of trichothiodystrophy (TTD), which is furthermore characterized by brittle hair and nails, ichthyosis and progressive mental and physical retardation (Hashimoto & Egly, 2009; Faghri *et al*, 2008). Besides XPB and XPD, only mutations in TTDA, the smallest subunit of TFIIH, have thus far been associated with photosensitive TTD (Giglia-Mari *et al*, 2004). Mutations in XPB, XPD and TTDA are rare and often found in compound heterozygous combinations. This, and the fact that so far only patients with mutations in three out of ten TFIIH subunits have been identified, reflects the fact that the TFIIH complex is essential for life (Park *et al*, 1992; De Boer *et al*, 1998; Theil *et al*, 2013; Andressoo *et al*, 2009).

In mice, disruption of XPD leads to embryonic lethality at the two-cell stage (De Boer *et al*, 1998). In contrast to other TFIIH subunits, yeast strains (Ranish *et al*, 2004) and mouse embryonic stem cells and fibroblasts (Theil *et al*, 2013) with complete inactivation of TTDA are viable, suggesting that TTDA is not essential for TFIIH basal transcription function. However, also disruption of TTDA leads to embryonic lethality in mice, albeit that embryos survive almost up to birth (Theil *et al*, 2013). Rare human TTD patients carry missense or nonsense mutations in *TTDA* that do not completely disrupt TTDA but lead to expression of a partially functional protein (Theil *et al*, 2011, 2013). This suggests that although TTDA appears to be less essential for transcription than other TFIIH subunits, full TTDA loss is not compatible with development of multicellular life (Theil *et al*, 2013; Moriwaki *et al*, 2014; Nonnekens *et al*, 2013). It is currently unclear why TTDA appears dispensable for transcription and viability of single cell systems but not of multicellular organisms and certain differentiated tissues.

The nematode *C. elegans* conserves many features of mammalian transcription and DNA repair (Bowman & Kelly, 2014; Boulton *et al*, 2002). We and others have previously shown that NER is conserved and protects genome integrity in germ cells and transcriptional integrity in somatic cells against DNA damage induced by UV light and other environmental and metabolic sources (Lans *et al*, 2010; Lans & Vermeulen, 2011, 2015; Astin *et al*, 2008; Rieckher *et al*, 2017; Meyer *et al*, 2007). Thus far, the composition and activity of *C. elegans* TFIIH has not been addressed. Here, we focus specifically on the *C. elegans* TTDA ortholog GTF-2H5 and show that this TFIIH factor is dispensable for development and viability under normal, unchallenged laboratory culture conditions. However, GTF-2H5 is essential for TFIIH stability and genome maintenance via NER and becomes vital to transcription when this is challenged.

## Results and discussion

### C. elegans lacking TTDA/GTF-2H5 is viable

To study the function of *TTDA/GTF2H5* in *C. elegans,* we characterized animals carrying a deletion mutation (*tm6360*) in the orthologous *gtf-2H5* gene. This gene only consists of two exons and is predicted to encode a 71 amino acid protein with an estimated molecular weight of 8.2 kD, which is similar as its human TTDA ortholog with which it shares 45% sequence identity. The *tm6360* allele was obtained from the National Bioresource Project for the nematode (Mitani, 2009) and represents a deletion of the entire second exon and flanking sequences (Fig 1A). By RT-PCR and RT-qPCR of the mRNA, we confirmed that *gtf-2H5* mutants do not express the second exon but found that the first exon is still expressed, albeit at >70% reduced levels (Fig 1B and C; Fig EV1A). To confirm whether indeed the *tm6360* allele encodes a truncated protein, we determined by PCR and sequencing of cDNA whether a mutated mRNA was being produced. In the *gtf-2H5* mutant, we detected by PCR a cDNA fragment stretching from the first exon of *gtf-2H5* (primer 1 in Fig 1A) to the last exon of the downstream *B0353.1* gene (primer 4 in Fig 1A; Fig EV1B), which after sequencing revealed that *gtf-2H5* mutants carrying the *tm6360* allele express low levels of a mutant mRNA consisting of the first *gtf-2H5* exon fused to the reverse sequence of the last exon of *B0353.1* (Fig EV1C). Gene prediction by FGENESH (Salamov & Solovyev, 2000) indicated that this mRNA encodes a mutated GTF-2H5 protein of which the C-terminal half is deleted and replaced by nonsense amino acid sequence (Fig 1D). Thus, *tm6360* is likely a strong loss-of-function allele.

**Figure 1.**
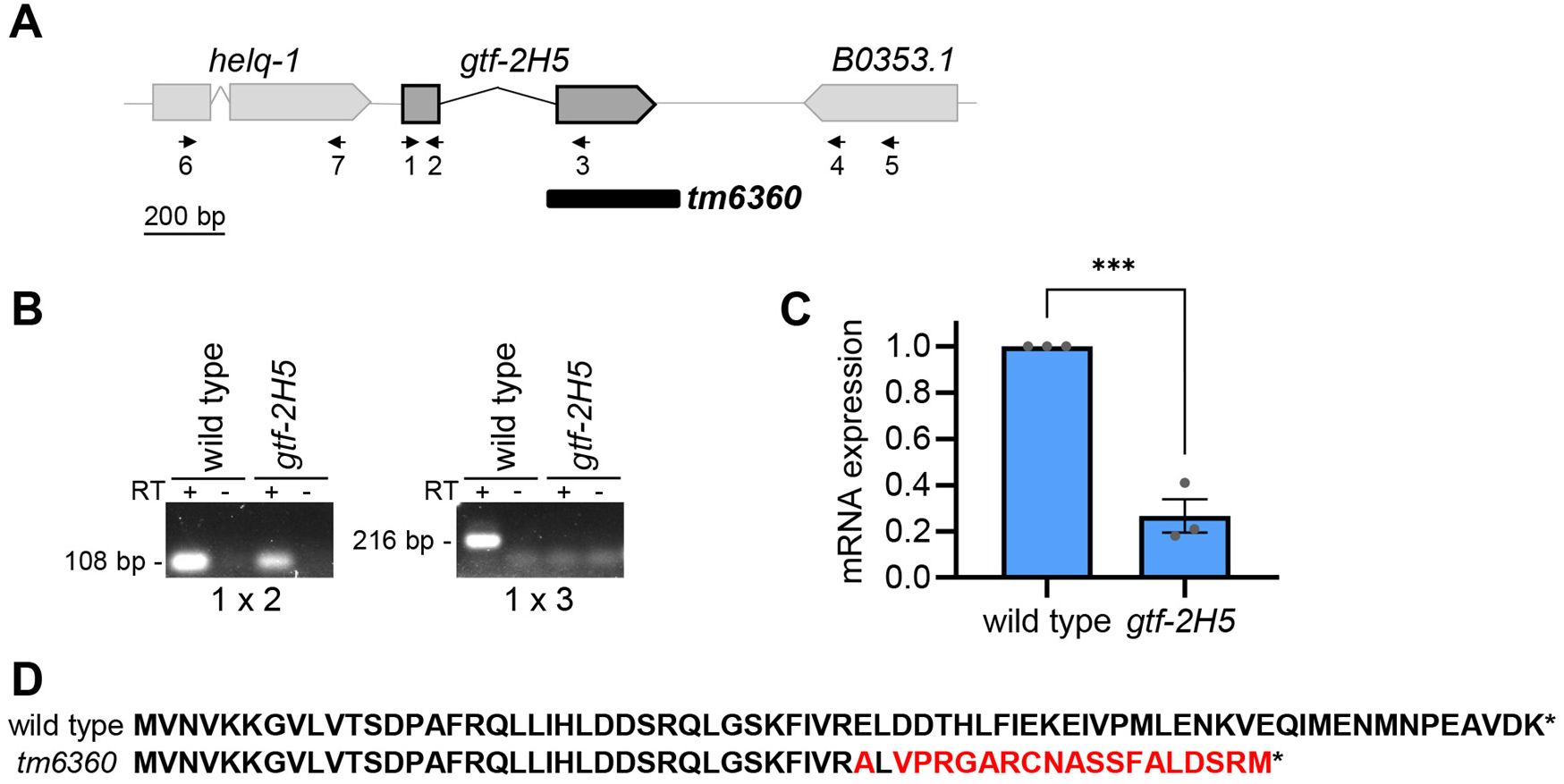
The *gtf-2H5(tm6360)* allele encodes a truncated protein. A. Schematic depiction of *gtf-2H5* locus with parts of flanking genes *helq-1* and *B0353.1*. Numbered arrows indicate positions of primers used in RT-(q)PCR. Primer sequences are listed in Table EV2. The deletion of the *tm6360* allele is indicated with a black box. B. PCR with primer pairs 1×2 or 1×3, as indicated in A, on cDNA of wild type or *gtf-2H5* animals. Lanes containing cDNA generated by reverse transcription of mRNA are labeled with ‘+’. As control, PCR on cDNA reaction samples without reverse transcriptase (RT) were included, labeled with ‘−‘. C. Relative mRNA expression levels of exon 1 of the *gtf-2H5* transcript as determined by qPCR on wild type and *gtf-2H5* mutant animals. mRNA levels were normalized to wild type. Results are plotted as average with SEM (error bars) of three independent experiments. Statistically significant difference (<0.001) is indicated by ***. D. Amino acid sequence of wild type GTF-2H5 and of the truncated protein predicted to be expressed by the *tm6360 gtf-2H5* allele. Red color indicate sequence that deviates from wild type.

TFIIH consists of ten subunits that are all conserved in *C. elegans* (Table 1) (Lans & Vermeulen, 2011) and which are all essential to life in mammals, including TTDA (Park *et al*, 1992; De Boer *et al*, 1998; Theil *et al*, 2013; Andressoo *et al*, 2009). Strikingly, we found that *gtf-2H5* mutants were viable, showed no embryonic lethality and produced similar brood size as wild type animals (Table 2). To test whether this might be due to the weakly expressed truncated GTF-2H5 protein being still partially functional, we examined whether full depletion of GTF-2H5 is lethal for *C. elegans.* To this end, we fused an auxin-inducible degradation (AID) tag (Zhang *et al*, 2015) together with GFP to GTF-2H5 by knocking in both tags at the C-terminus of the *gtf-2H5* gene using CRISPR/Cas9 (Fig 2A). As comparison, we also knocked in AID and GFP at the N-terminus of *gtf-2H1* (Fig 2B), the *C. elegans* ortholog of TFIIH subunit GTF2H1/p62, which in humans and yeast is essential for TFIIH integrity and function (Luo *et al*, 2015; Ribeiro-Silva *et al*, 2018; Barnett *et al*, 2020; Fischer *et al*, 1992). For simplicity, we will refer to the AID::GFP tag with ‘AG’ and to these knockin animals as *gtf-2H5::AG and AG::gtf-2H1*. Both knockin animals were fully viable and similarly expressed AG-tagged TFIIH in nuclei of all tissues (Fig 2C; Fig EV1D). Also, both transgenic animals displayed normal, wild type UV survival in an assay that measures TC-NER (Fig EV2A) (Lans *et al*, 2010; van der Woude & Lans, 2021), indicating that the TFIIH complex is intact and functional when either GTF-2H5 or GTF-2H1 is tagged. We crossed both strains with animals expressing Arabidopsis TIR1 (fused to mRuby) specifically in germ cells and early embryos under control of the *sun-1* promoter (Zhang *et al*, 2015; Minn *et al*, 2009). Exposure to auxin, which activates the TIR1 E3 ubiquitin ligase complex that ubiquitylates the AID tag, led to full depletion of both fusion proteins in the germline (Fig EV2B). However, only the depletion of AID-tagged GTF-2H1, and not that of GTF-2H5, caused complete embryonic lethality (Fig 2D and E). Previously, we also showed that auxin-induced depletion of the TFIIH subunit XPB-1 causes developmental arrest (Sabatella *et al*, 2021). As animals with truncated or depleted GTF-2H5 do not show any embryonic lethality, we conclude that unlike other TFIIH subunits, GTF-2H5 is not essential for embryogenesis and viability in *C. elegans.*

**Table 1.** *C. elegans* TFIIH subunits.

| ***C. elegans* gene** | **human ortholog** |
| --- | --- |
| *xpb-1* | *ERCC3 (XPB)* |
| *xpd-1* | *ERCC2 (XPD)* |
| *gtf-2H1* | *GTF2H1 (p62)* |
| *gtf-2H2C* | *GTF2H2 (p44)* |
| *gtf-2H3* | *GTF2H3 (p34)* |
| *gtf-2H4* | *GTF2H4 (p52)* |
| *gtf-2H5* | *GTF2H5 (p8)* |
| *mnat-1* | *MNAT1 (MAT1)* |
| *cyh-3* | *CCNH (Cyclin H)* |
| *cdk-7* | *CDK7* |

**Table 2.** Brood size of wild type and *gtf-2H5* animals.

| **strain** | **brood size** |
| --- | --- |
|  | **average±sem (n)** |
| wild type | 215±14 (n=20) |
| *gtf-2H5(tm6360)* | 212±8 (n=20) |

**Figure 2.**
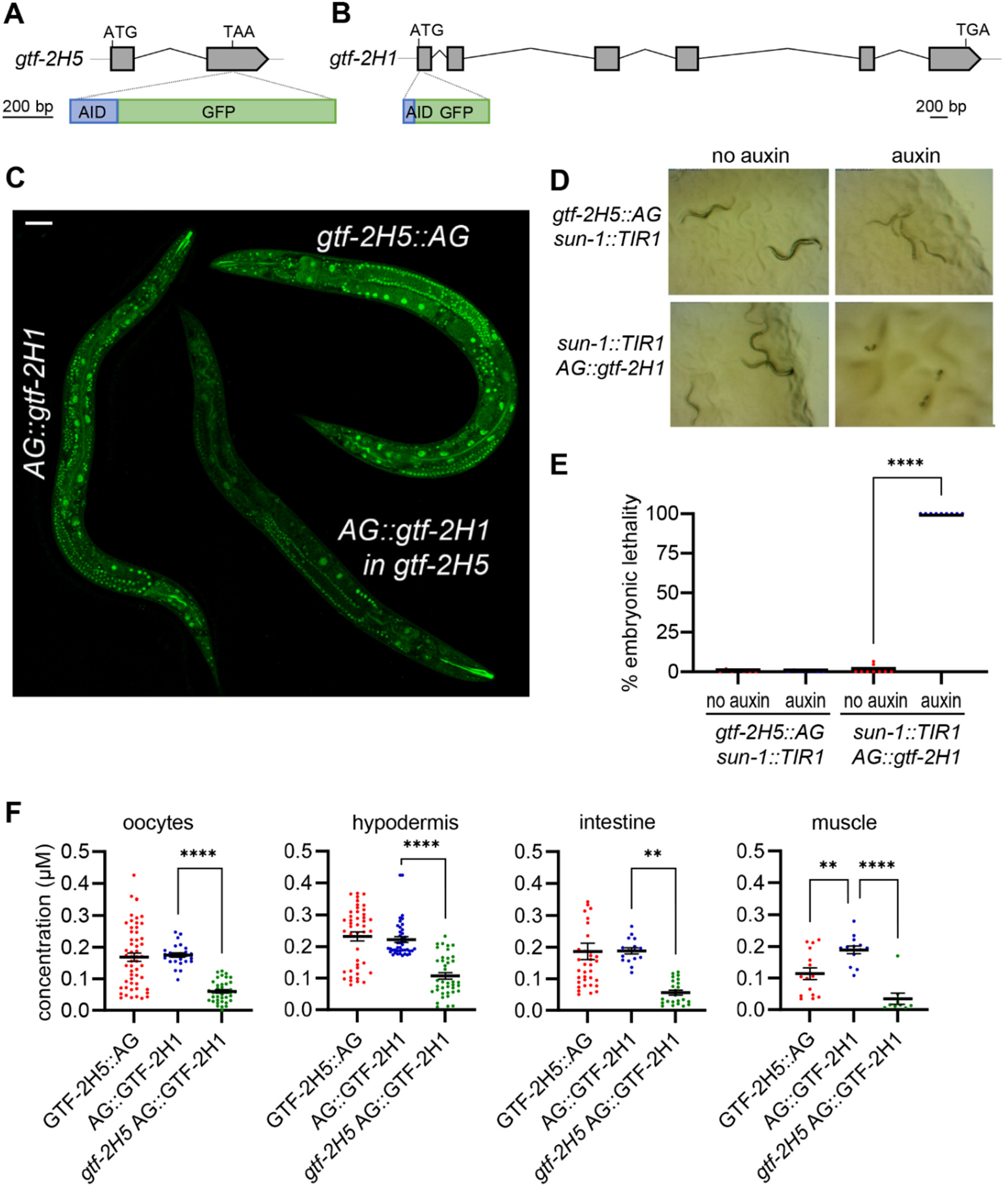
GTF-2H5 and GTF-2H1 are ubiquitously expressed but at different concentrations. A. Schematic depiction of the *gtf-2H5* locus and C-terminal knockin site of the AID::GFP tag. B. Schematic depiction of the *gtf-2H1* locus and N-terminal knockin site of the AID::GFP tag. C. Composite overview image generated by merging independent confocal scans of *gtf-2H5::AG, AG::gtf-2H1* and *gtf-2H5; AG::gtf-2H1*. Scale bar: 50 μm. D. Stereo microscope view of the offspring of *gtf-2H5::AG* and *AG::gtf-2H1* animals expressing TIR1 under control of the *sun-1* promoter grown in absence or presence of 1 mM auxin. Only auxin-induced depletion of *AG::gtf-2H1* leads to embryonic lethality. E. Quantification of the experiment described in D. Shown is a scatter dot plot of the average percentage embryonic lethality observed on at least seven plates for each condition in two independent experiments. Statistically significant difference (<0.0001) is indicated by ****. F. Quantification of GTF-2H5::AG and AG::GTF-2H1 concentration in nuclei of oocytes, hypodermal, intestinal and muscles cells of wild type and *gtf-2H5* animals. Concentration was determined by comparison of the average fluorescence levels in the entire nucleus to the fluorescence of known concentrations of purified GFP.

### GTF-2H5 promotes TFIIH stability in multiple tissues in vivo

Both GTF-2H5 as well as GTF-2H1 showed exclusive nuclear expression in various tissues of *C. elegans* (Fig EV1D). To have an idea of the concentration of TFIIH molecules present *in vivo* in nuclei of different tissues, we determined GTF-2H5::AG and AG::GTF-2H1 concentrations in several readily visible tissues by carefully comparing average nuclear fluorescence signals to that of known GFP concentrations. This showed comparable concentrations of around 0.2 μM for GTF-2H1 in oocyte, hypodermal, intestinal and muscle nuclei (Fig 2F), which is strikingly similar to concentration estimations made of Pol II in human fibroblasts (0.18 μM) using a similar method (Steurer *et al*, 2018). GTF-2H5 concentrations were mostly similar, but showed higher variation between individual nuclei and a lower concentration in muscle cells. This may reflect the notion that this factor can dynamically associate with TFIIH (Giglia-Mari *et al*, 2006). We quantified TFIIH concentration in brightly fluorescent cells that were easily discernable, but noticed that in many other nuclei, such as for instance in head neurons, expression levels seemed much lower (Fig EV1D). As nuclei in neurons are smaller than those in oocytes, hypodermal and intestinal cells, the TFIIH concentration will also be lower in these cell types. Similar analysis of mouse XPB-YFP fluorescence levels in tissues of XPB-YFP knockin mice also showed that TFIIH levels vary depending on the cell type, which, interestingly, was found to correlate to transcriptional activity in these cells (Donnio *et al*, 2019). Thus, possibly also in *C. elegans* TFIIH levels can be used as quantitative biomarker of transcriptional activity. This correlation, however, is difficult to prove as quantification of nascent mRNA by 5-ethynyl uridine labeling in different tissues of *C. elegans* does not produce sufficiently reproducible results in our hands (Sabatella *et al*, 2021).

To determine the impact of GTF-2H5 loss on TFIIH levels, we crossed *AG::gft-2H1* animals with *gtf-2H5* mutants and found that loss of GTF-2H5 led to reduced levels of AG::GTF2H1 in all tested tissues by more than 50% (Fig 2F). These results are in line with lowered TFIIH protein levels observed in patient and mouse fibroblasts (Theil *et al*, 2013; Botta *et al*, 2002), while TFIIH subunit mRNA levels are unaffected (Vermeulen *et al*, 2000), and indicate that GTF-2H5 has an evolutionary conserved function in maintaining steady state levels of TFIIH, not only *in vitro* but also in various tissues *in vivo*.

### GTF-2H5 is essential for nucleotide excision repair

Because of the lowered TFIIH levels, we tested whether specific functions of TFIIH were affected and first focused on its role in NER. We and others have previously shown that UV-induced DNA damage causes embryonic lethality in the absence of GG-NER (Lans *et al*, 2010; Lans & Vermeulen, 2015; Astin *et al*, 2008; van der Woude & Lans, 2021). We observed that *gtf-2H5* mutants are as sensitive to UV-induced DNA damage as NER-deficient *xpa-1* mutants, showing severe embryonic lethality upon UV irradiation (Fig 3A). These results indicate that the *tm6360* allele is a null allele with respect to GTF-2H5 function in NER.

**Figure 3.**
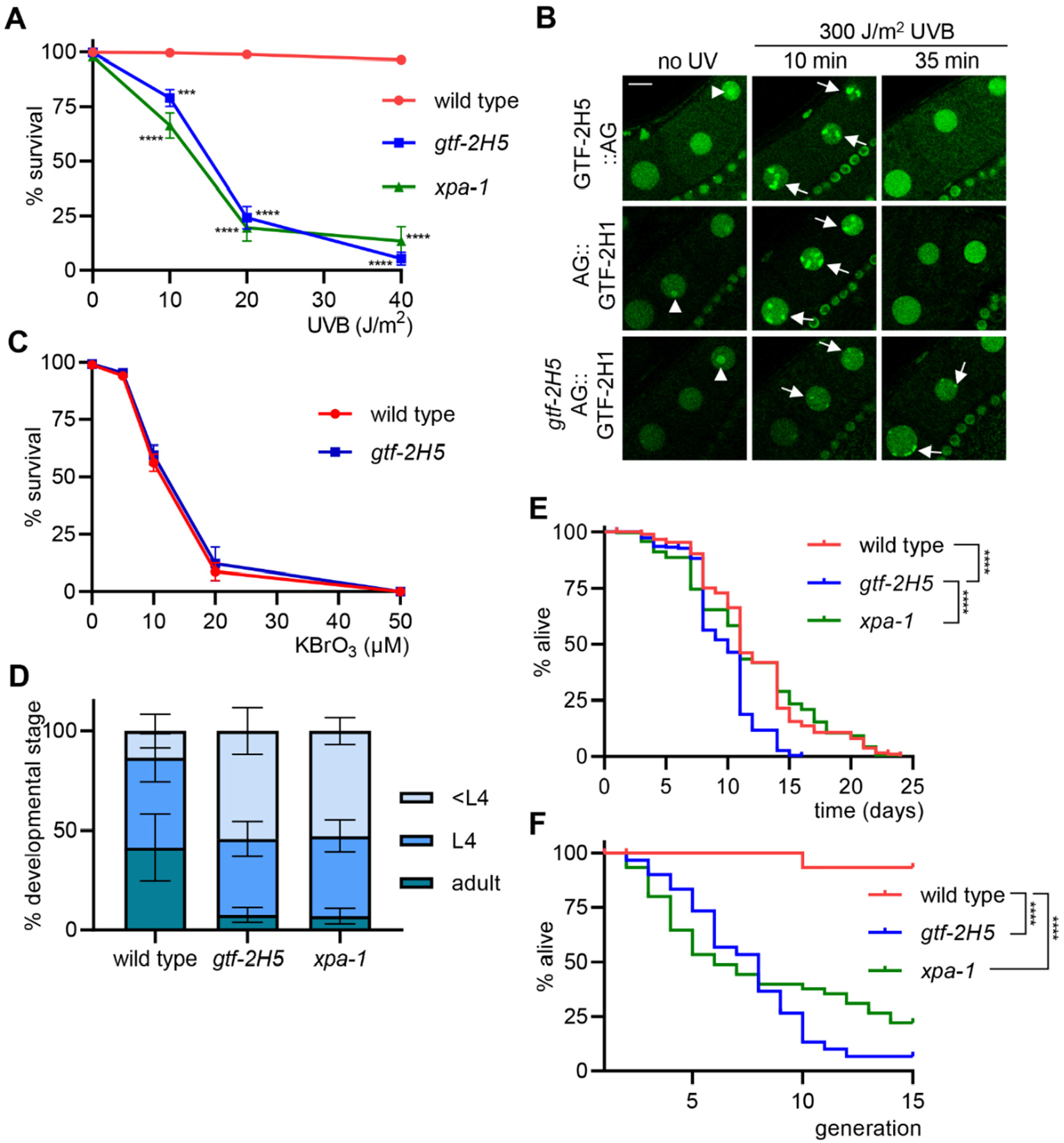
*gtf-2H5* animals are NER deficient and show diminished growth and lifespan. A. Germ cell and embryo survival assay after UVB irradiation of germ cells in young adult wild type, *gtf-2H5* and *xpa-1* animals. The percentages of hatched eggs (survival) after UVB irradiation are plotted against the applied UV-B doses. Results are plotted as average with SEM (error bars) of at least six experiments. Statistically significant difference compared to wild type for each dose is indicated by *** (<0.001) or **** (<0.0001). B. Representative images showing real-time recruitment of GTF-2H5::AG or AG::GTF-2H1 to UVB-damaged chromosomes in oocytes of living wild type (upper and middle panel) or *gtf-2H5* (lower panel) animals, before UVB irradiation (no UV) and 10 min and 35 min after 300 J/m^2^ UVB irradiation. Recruitment to paired homologous chromosomes are indicated with arrows. Nucleoli are indicated with an arrowhead. Scale bar: 10 μm C. Survival assay after incubation of wild type and *gtf-2H5* L1/L2 larvae with increasing concentrations of KBrO_3_ for 24 h, which induces oxidative DNA damage. The percentages of non-arrested, developing larvae (survival) are plotted against the applied KBrO_3_ concentration. Results are plotted as average with SEM (error bars) of two independent experiments each performed in triplicate. D. Quantification of larval growth of wild type, *gtf-2H5* and *xpa-1* animals by determining the percentage of adult, L4 and younger than L4 (<L4) animals observed 48 h after animals are laid as eggs at 25°C. E. Post-mitotic lifespan analysis showing the percentage of alive adult wild type (n=290), *gtf-2H5* (n=300) and *xpa-1* (n=285) animals per day. Statistically significant difference (<0.0001) is indicated by ****. F. Replicative lifespan analysis showing the percentage survival of successive generations of wild type, *gtf-2H5* and *xpa-1* animals if, in each generation, one animal is passaged. Depicted are cumulative results from at least two independent experiments (n=15 per experiment). Statistically significant difference (<0.0001) is indicated by ****.

To gain further insight into the cause of UV hypersensitivity of *gtf-2H5* mutant animals, we visualized recruitment of TFIIH to DNA damage. In *C. elegans* oocytes, DNA is condensed and organized into six pairs of bivalents, i.e., paired homologous chromosomes, that are readily discernable by microscopy e.g. when stained with DAPI (Fig EV2C, left panel). We showed previously that upon UV irradiation, the NER endonuclease ERCC-1/XPF-1 rapidly re-localizes from the nucleoplasm to damaged chromosomes, reflecting its activity in NER (Sabatella *et al*, 2021). This damaged-DNA binding is dependent on GG-NER and lasts for approximately half an hour until UV photolesions are repaired and NER is completed. In line with this, we observed clear re-localization of both GTF-2H5::AG and AG::GTF-2H1 to oocyte bivalents 10 min after UV irradiation, while imaging living (Fig 3B) or fixed (Fig EV2C) animals, reflecting TFIIH binding to UV-damaged DNA. This recruitment was not observed anymore 35 min after UV, indicating completion of repair, as previously shown (Sabatella *et al*, 2021). Next, we tested re-localization of AG::GTF-2H1 in *gtf-2H5* mutants and strikingly observed that its UV-induced recruitment to damaged chromosomes was strongly reduced after 10 min and persisted after 35 min (Fig 3B). We observed similar *gtf-2H5-*dependent recruitment to damaged bivalents of another TFIIH subunit, XPB-1 (Fig EV2D), of which we previously had generated AID::GFP knockin animals (Sabatella *et al*, 2021). Together with the strong UV hypersensitivity of *gtf-2H5* mutants, these results indicate that GTF-2H5 is needed for efficient binding of TFIIH to damaged DNA and that in absence of GTF-2H5 repair cannot be completed. Also *in vitro* and in cultured mammalian cells TTDA was found to be essential for NER and the recruitment of TFIIH to sites of UV-induced DNA damage, likely by stimulating XPB ATPase activity and TFIIH translocase activity together with XPC and XPA (Theil *et al*, 2013; Giglia-Mari *et al*, 2004; Coin *et al*, 2006; Kappenberger *et al*, 2021). Our results show that GTF-2H5 has a similar, evolutionary conserved role in stimulating TFIIH activity in NER in oocytes of *C. elegans.* Besides NER, TTDA has been implicated in repair of oxidative DNA damage, as TTDA knockout mouse embryonic stem cells are sensitive to ionizing radiation and KBrO_3_ (Theil *et al*, 2013). However, when comparing KBrO_3_ sensitivity of *C. elegans gtf-2H5* mutants and wild type animals, we did not observe any obvious hypersensitivity (Fig 3C).

Since *gtf-2H5* mutants are as UV hypersensitive as *xpa-1* mutants, we tested whether these animals also exhibit phenotypes that we previously reported to be caused by accumulating DNA damage and mutations due to NER deficiency. A significant proportion of *C. elegans* NER deficient animals in a population exhibit growth delay due to spontaneous DNA damage (Lans *et al*, 2013). Indeed, we observed that a percentage of *gtf-2H5* animals exhibited delayed development, just as *xpa-1* animals, in comparison to wild type (Fig 3D). Also, we previously showed that NER mutants do not have reduced postmitotic lifespan, i.e. longevity measured from the onset of adulthood, but do show severely reduced replicative lifespan, i.e. transgenerational replicative capacity (Lans *et al*, 2013). We confirmed again that *xpa-1* mutants have a postmitotic lifespan comparable to wild type animals (Fig 3E), but that their ability to proliferate and reproduce over generations strongly declines (Fig 3F). To our surprise, however, we found that *gtf-2H5* mutants show both a shortened postmitotic (Fig 3E) and replicative (Fig 3F) lifespan, which appear both more reduced than of *xpa-1* mutants. Thus, besides being clearly NER defective, *gtf-2H5* animals likely carry a defect in an additional biologic pathway that causes them to live shorter.

### GTF-2H5 is required for transcription when this is challenged

TTD patients with mutations in TTDA, XPB and XPD exhibit typical TTD features like brittle hair and nails and ichthyosis and, in addition, photosensitivity and progressive neuropathy that are reminiscent of xeroderma pigmentosum (Hashimoto & Egly, 2009; Faghri et al, 2008). Photosensitivity and part of the neuropathy are considered to be due to defects in NER, but the other features are thought to be caused by limited availability of TFIIH, due to its instability and lowered levels that lead to its exhaustion in terminally differentiated cells, such as skin and hair keratinocytes, before the transcriptional program in these cells has ended (De Boer et al, 1998b; Theil et al, 2014). As TFIIH is needed for transcription of abundant proteins in these cells, such as cysteine-rich matrix proteins, reduced amounts of TFIIH likely explain the sulfur-deficient brittle hair and nails and scaly skin features. Thus, we postulate that, similarly, the lowered TFIIH levels in *gtf-2H5* animals lead to TFIIH exhaustion and limited TFIIH availability for transcription initiation in older animals, causing cell dysfunction and reduced lifespan.

To test whether the lowered TFIIH amounts in *gtf-2H5* animals indeed become restrictive in circumstances with altered transcriptional demand, we tested whether a transcription defect might become apparent in these animals when transcription is challenged. As transcription cannot be reliably measured with 5-ethynyl uridine labeling, we crossed *gtf-2H5* mutants with animals expressing GFP in body muscle cells under control of the *eft-3* promoter (Zhang *et al*, 2015), and quantified GFP fluorescence levels as proxy for transcriptional competence (Sabatella *et al*, 2021). This showed no difference between wild type and *gtf-2H5* animals (Fig 4A and B), indicating that in unperturbed conditions *gtf-2H5* animals are fully transcriptionally competent for this muscle-specific transgene.

**Figure 4.**
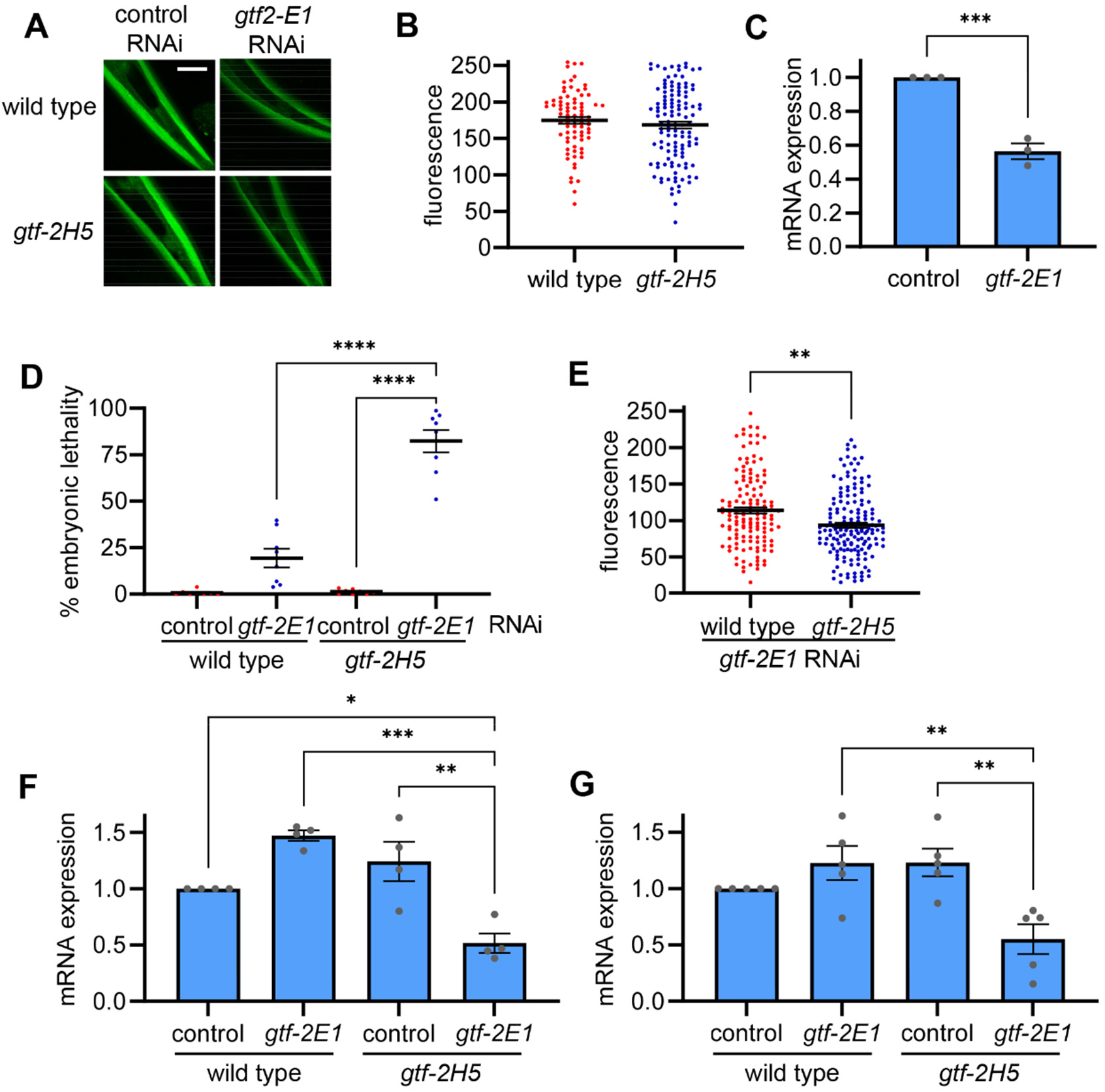
GTF-2H5 promotes transcription when this is challenged. A) Representative images of wild type and *gtf-2H5* animals expressing AID::GFP under control of *eft-3* promoter in body wall muscles, shown here in the head of *C. elegans,* grown on control or *gtf-2E1* RNAi. Scale bar: 25 μm B) Scatter dot plot showing average and SEM of the relative GFP fluorescence levels in head muscle cells of wild-type and *gtf-2H5* animals, as depicted in A. C) Relative *gtf-2E1* mRNA levels as determined by qPCR on animals grown on control or *gtf-2E1* RNAi. Results are normalized to control RNAi and plotted as average with SEM (error bars) of three independent experiments. Statistically significant difference (<0.001) is indicated by ***. D) Scatter dot plot showing the average percentage with SEM of embryonic lethality observed on eight plates in two independent experiments with either wild type animals or *gtf-2H5* animals (both also carrying the *eft-3::GFP* transgene used in A and B, grown on control or *gtf-2E1* RNAi food. Statistically significant differences are indicated by **** (<0.0001). E) Scatter dot plot showing average and SEM of the relative GFP fluorescence levels in head muscle cells of wild-type and *gtf-2H5* animals grown on *gtf-2E1* RNAi food, as depicted in A. Statistically significant differences are indicated by ** (<0.01). F) Relative *cdc-42* mRNA levels as determined by qPCR on wild type or *gtf-2H5* animals grown on control or *gtf-2E1* RNAi. Results are normalized to wild type on control RNAi and plotted as average with SEM (error bars) of four independent experiments. Statistically significant difference is indicated by *(<0.05), ** (<0.01) or *** (<0.001). G) Relative *pmp-3* mRNA levels as determined by qPCR on wild type or *gtf-2H5* animals grown on control or *gtf-2E1* RNAi. Results are normalized to wild type on control RNAi and plotted as average with SEM (error bars) of five independent experiments. Statistically significant differences are indicated by ** (<0.01).

Next, we compromised transcription by limiting the availability of another transcription initiation factor, without compromising NER. To this end, we cultured wild type and *gtf-2H5* animals on *gtf-2E1* RNAi bacteria to diminish protein levels of the essential TFIIE transcription factor complex, which stimulates TFIIH activity and helps anchor it within the Pol II pre-incision complex, but has no role in NER (Yamamoto *et al*, 2001; Compe *et al*, 2019; Ohkuma & Roeder, 1994; Theil *et al*, 2017; Kuschal *et al*, 2016; Park *et al*, 1995). *gtf-2E1* RNAi food is not fully efficient and causes only partial knockdown of *gtf-2E1* transcripts and therefore only mild embryonic lethality in wild type animals (Fig 4C and D). Strikingly, however, *gtf-2H5* animals grown on the same *gtf-2E1* RNAi food showed very high levels of embryonic lethality. By measuring GFP fluorescence in body wall muscles, we observed that partial *gtf-2E1* knockdown lowered GFP proteins levels in wild type animals, which was further enhanced in *gtf-2H5* animals (Fig 4A and E). Thus, in conditions of limited availability of the TFIIE transcription factor and lowered gene expression, GTF-2H5 becomes essential for transcription efficiency and viability. To confirm this in another way, we measured the transcript levels of two housekeeping genes, *cdc-42* and *pmp-3*, by qPCR in cDNA generated from RNA isolated from the same amount of non-gravid adult wild type or *gtf-2H5* animals grown on control or *gtf-2E1* RNAi. This showed that *gft-2H5* knockout or *gtf2E1* knockdown alone did not result in lowered transcript levels, but the combined loss of *gtf-2H5* and *gtf-2E1* strongly reduced transcription of both housekeeping genes (Fig 4F and G). These results therefore suggest that in *C. elegans* GTF-2H5 (and thus high steady-state TFIIH levels) is dispensable for transcription throughout most of *C. elegans* lifespan in unperturbed laboratory culturing conditions, but becomes essential for transcription if this process is somehow compromised or challenged.

This notion may explain contradicting results in previous studies of TTDA’s importance to transcription. Initially, upon its identification, human TTDA was considered a NER-specific TFIIH factor dispensable for transcription. TTD-A patient cells had no obvious transcriptional defects and purified TTDA did not stimulate TFIIH-dependent transcription *in vitro* (Vermeulen *et al*, 2000; Giglia-Mari *et al*, 2006; Coin *et al*, 2006; Kainov *et al*, 2008). However, TTDA knockout mouse embryonic stem cells did show reduced transcription levels (Theil *et al*, 2013) and a reconstituted transcription assay using different recombinant TFIIH also showed that transcription was stimulated by TTDA (Aguilar-Fuentes *et al*, 2008). Furthermore, yeast lacking TTDA ortholog TFB5 did not show any major change in gene expression as observed by microarray, but still TFB5 was found to stimulate transcription *in vitro* and efficient transcription *in vivo* when there was high demand such as in changing environmental conditions (Ranish *et al*, 2004). Thus, it appears that depending on the particular conditions of the assay or cell type used, contingent on transcriptional demand, the absence of TTDA/GTF-2H5 may or may not be a limiting factor. This feature may explain why cultured yeast and mammalian cells and *C. elegans* can thrive without TTDA/GTF-2H5, whereas mice with full TTDA loss are not viable, likely because mammalian embryogenesis requires high transcriptional capacity (Theil *et al*, 2014).

High resolution cryo-EM structures of the human preinitiation complex do not suggest a direct role for TTDA in loading or DNA interaction of TFIIH at promoters (Aibara *et al*, 2021). Therefore, to test whether the importance of GTF-2H5 for compromised transcription is due to its ability of stabilizing TFIIH, we tested if lowering TFIIH levels in another way, by partially depleting the TFIIH subunit GTF-2H2C (p44/GTF2H2 in humans; Table 1), led to similar enhanced embryonic lethality when transcription is compromised. To this end, we cultured wild type animals on an equal mixture of control RNAi and *gtf-2H2C* RNAi food, which destabilized TFIIH, as visualized by lowered AG::GTF-2H1 levels (Fig 5A), and caused incompletely penetrant embryonic lethality (Fig 5B). However, when *gtf-2H2C* RNAi food was equally mixed with *gtf-2E1* RNAi food, this led to very high levels of embryonic lethality. These results confirm that unstable and lowered levels of TFIIH compromise transcription when this is challenged.

**Figure 5.**
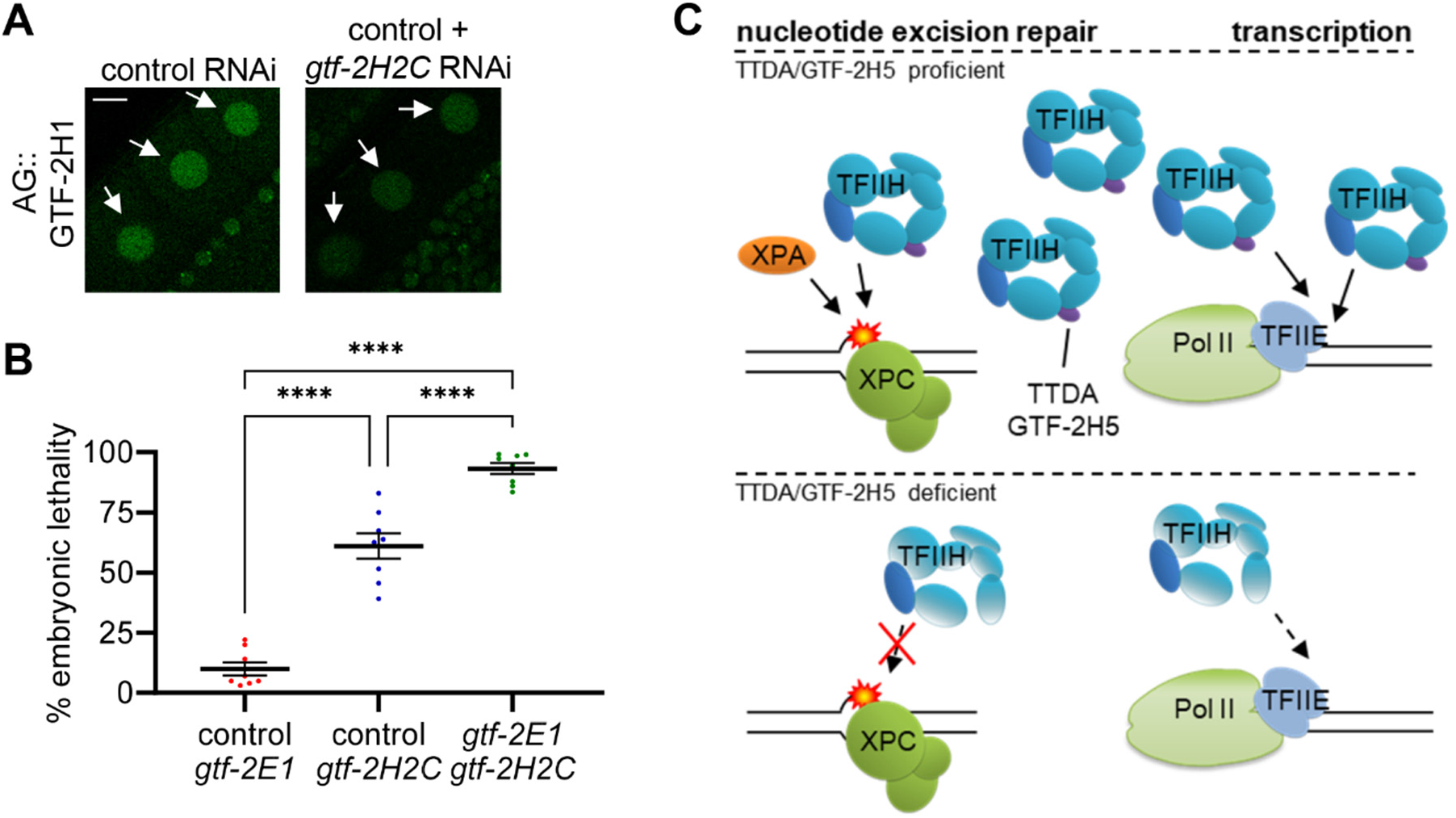
Synthetic lethality between GTF-2H2C and GTF-2E1 and model for GTF-2H5 activity. A) Representative images of AG::GTF-2H1 fluorescence in oocyte nuclei (indicated by arrows) of *AG::gtf-2H1* knockin animals grown on control RNAi or a 1:1 mixture of control and *gtf-2H2C* RNAi food. Scale bar: 10 μm B) Scatter dot plot showing the average percentage with SEM of embryonic lethality observed on eight plates in two independent experiments with wild type animals grown on 1:1 mixtures of control RNAi with *gtf-2E1* RNAi food, control RNAi with *gtf-2H2C* RNAi food or *gtf-2E1* RNAi and *gtf-2H2C* RNAi food. C) Model for TTDA/GTF-2H5 involvement in nucleotide excision repair and transcription initiation. In wild type cells (upper part), TTDA/GTF-2H5 is the smallest subunit of a fully functional TFIIH complex, which exists in sufficiently high concentrations to promote nucleotide excision repair together with XPC and XPA and transcription initiation by RNA polymerase II (Pol II) together with TFIIE (and other not depicted repair and transcription factors). In TTDA/GTF-2H5 deficient cells (lower part), the TFIIH complex is not efficiently recruited and active in nucleotide excision repair. Also, as the complex is unstable and exists in low concentrations, it can only support transcription initiation if this is not too demanding.

### gtf-2H5 C. elegans mutants as model for trichothiodystrophy

TFIIH is an essential transcription and DNA repair factor. While thus far all TFIIH subunits were thought to be essential to multicellular life, including TTDA/GTF2H5, here we show that *C. elegans* strains with truncated or complete depletion of GTF-2H5 are viable. As *gtf-2H5* mutants are strongly UV-hypersensitive, similar as *xpa-1* loss-of-function mutants, and show impaired recruitment of TFIIH to UV-induced DNA damage, GTF-2H5 is an essential NER factor (Fig 5C). With the characterization of GTF-2H5 presented in this study, and with it of the entire TFIIH complex, we and others have shown full functional conservation of both the GG-and TC-NER pathway in *C. elegans* (Astin *et al*, 2008; Babu & Schumacher, 2016; Babu *et al*, 2014; Lans *et al*, 2010, 2013; Lans & Vermeulen, 2011; Geijer *et al*, 2021; Lee *et al*, 2002; Park *et al*, 2004; Stergiou *et al*, 2007; Sabatella *et al*, 2021; Hartman *et al*, 1989; Meyer *et al*, 2007).

In recent years, besides mutations in XPB, XPD and TTDA, multiple additional mutations in genes have been identified to cause TTD or TTD-like syndromes, among which are mutations in transcription initiation factor TFIIE, which all affect the stability of proteins involved in different steps of transcription and translation (Kuschal *et al*, 2016; Theil *et al*, 2017; Botta *et al*, 2021; Kuo *et al*, 2019; Theil *et al*, 2019; Tessarech *et al*, 2020; Corbett *et al*, 2015). Together, these have put forward the unifying idea that TTD is caused by instability of proteins involved in gene expression. Our results are in line with this idea and confirm that loss of GTF-2H5 leads to instability of the transcription initiation factor TFIIH, which may lead to transcription problems in cells of aged animals due to TFIIH exhaustion (Fig 5C). Furthermore, we find that when transcription initiation is hampered, by lowering of TFIIE levels, the reduced TFIIH levels in *gtf-2H5* mutants become restrictive and cause strong embryonic lethality. Thus, in contrast to its function in NER, TTDA appears mostly dispensable for TFIIH function in facilitating transcription initiation by RNA polymerase I and II. However, in conditions of transcription stress, when there is demand for high TFIIH concentrations due to e.g high transcriptional output or reduced availability of other transcription initiation factors, the TFIIH-stabilizing ability of TTDA becomes essential to ensure sufficiently high TFIIH levels for transcription to initiate. These findings indicate that *gtf-2H5* mutant *C. elegans* can be an advantageous multicellular animal model for studying genetic and developmental pathogenic aspects of TTD features.

## Material and methods

### C. elegans strains and culture

*C. elegans* strains used are listed in Table EV1. *C. elegans* was cultured according to standard methods on nematode growth media (NGM) agar plates seeded with *Escherichia coli* OP50. All mutants were backcrossed against wild type strain, which was Bristol N2. *gtf-2H5::AG* knockin animals were generated using gRNA with targeting sequence CATGGAAAATATGAATCCGG and as homology directed repair template a heteroduplex DNA fragment generated by mixing PCR products (Dokshin *et al*, 2018) obtained with primers combinations 5’-GAACTCGACGATACGCATTTG-3’/5’-GCCTAAAACATGAAGCCTGTTG-3’ and 5’-GTATCCTTCAATCCACCGTTC-3’/5’-TTTGTATAGTTCGTCCATGCC-3’ on a gene fragment consisting of *AID::GFP* sequences flanked by 200 bp homology arms from the *gtf-2H5* locus. *AG::gtf-2H1* knockin animals were generated using gRNA with targeting sequence TTTTCAGATGTCAGACGAGT and as homology directed repair template a gene fragment consisting of *AID::GFP* sequences flanked by 100 bp homology arms from the *gtf-2H1* locus in pUCIDT-KANA (Integrated DNA technologies). sgRNA and plasmids were injected in the Cas9dPiRNA expressing strain HCL67 (a kind gift from Heng-Chi Lee) (Zhang *et al*, 2018). Knockin animals were verified by genotyping PCR and sequencing, after which the Cas9 was removed by backcrossing against wild type. RNAi bacteria were obtained from the *Caenorhabditis elegans* RNAi feeding library (Kamath *et al*, 2003). Control RNAi was vector pPD129.36 (a gift from Andrew Fire).

### Brood size, growth and embryonic lethality

To measure brood size, L4 animals were individually seeded and transferred to a new culture plate every day, after which eggs laid were counted. To measure growth, adult animals were allowed to lay eggs for 4 h. After 48 h at 25°C, the growth of the progeny was scored by counting the amount of adult, L4 and younger than L4 animals. To measure embryonic lethality of AID-tagged transgenic strains, late L4 animals were mock treated or exposed to 1 mM auxin (3-indoleacetic acid, Sigma) for 24 h. To measure embryonic lethality of RNAi-fed strains, animals were first grown for one generation on RNAi food. Next, four animals per plate were allowed to lay eggs for 3 h, after which hatched and unhatched eggs were scored.

### Lifespan assays

To measure post-mitotic lifespan, standard longevity assays were performed at 20°C as previously described with day 1 defined as the day when animals reached adulthood (Lans & Jansen, 2007). Animals were scored every 1 to 3 days. To measure transgenerational replicative lifespan, assays were performed as previously described (Lans *et al*, 2013). In brief, 15 animals per strain were individually seeded and cultured to produce progeny at 25°C. From each progeny, a single animal was randomly picked, transferred to a fresh culture plate and grown to produce new offspring, which was repeated for 15 generations. Progeny was considered to have lost viability if animals arrested development, produced no or inviable progeny or died before reproduction. Animals that crawled off the plate were censored.

### RT-PCR and RT-qPCR

For RT-PCR, animals were lysed in TRIzol (Qiagen) and RNA was purified using RNeasy spin columns (Qiagen). cDNA was generated using Superscript II Reverse Transcriptase (Invitrogen) according to manufacturer’s instructions. For RT-qPCR, RNA was isolated and cDNA generated from 10 young adult animals per strain and condition using the Power SYBR Green Cells-to-Ct kit (Invitrogen) according to the method described by Chauve *et al* (Chauve *et al*, 2020). qPCR was performed using PowerUp SYBR Green Master Mix (Invitrogen), according to manufacturer’s instructions, with 58°C as annealing temperature and 1 min elongation time. Primers used for RT-PCR and RT-qPCR are listed in Table EV2. *cdc-42* and *pmp-3* cDNA were used as reference genes, unless otherwise stated.

### Imaging and GTF-2H5 and GTF-2H1 concentration measurements

For microscopy of living animals, animals were mounted on 2% agar pads in M9 containing 10 mM NaN3 (Sigma) and imaged on a Zeiss LSM700 or Leica SP8 confocal microscope. For microscopy of fixed animals, animals were fixed on Poly-L-lysine hydrobromide (Sigma) slides with 4% paraformaldehyde in PBS and slides were mounted using Vectashield with DAPI (Vector laboratories). Fluorescence intensities of AID::GFP driven by *eft-3* promoter in head body wall muscles were imaged and quantified using ImageJ software. To estimate concentrations of GTF-2H5::AG and AG::GTF-2H1, we imaged GFP fluorescence intensity of Z-slices of individual nuclei of the indicated cell-types. Average fluorescence intensities of each nucleus were then calibrated to fluorescence intensities of different concentrations of purified eGFP (Biovision), using similar imaging condition, to derive the protein concentration.

### Survival assays

UV survival was carried out as described (Lans *et al*, 2010; van der Woude & Lans, 2021). For UV irradiation, Philips TL-12 UV-B tubes (40W) were used. To determine UV-induced embryonic lethality in Fig 3A, staged young adult worms were washed, UVB-irradiated on agar plates without food and allowed to recover for 24 h on plates with OP50 bacteria. Next, five adults were allowed to lay eggs for 3 h on 6 cm plates seeded with HT115 bacteria, in five-fold for each UVB dose and 24 h later, the amount of hatched and unhatched (dead) eggs was counted. To determine UV-induced larval growth arrest in Fig EV2A, eggs were collected by hypochlorite treatment of adult animals and plated onto agar plates with HT115 bacteria. After ~16 h, L1 larvae were UVB-irradiated and allowed to recover for 48 h. Survival was determined by counting animals arrested at the L1/L2 stages and animals that developed beyond the L2 stage. To determine KBrO_3_ sensitivity, eggs were collected by hypochlorite treatment and allowed to hatch in S basal medium containing OP50 bacteria for 16 h at 21°C while shaking. Next, the indicated KBrO_3_ concentrations were added to the liquid cultures, in triplicate. After 24 h incubation, animals were plated on 6 cm culture plates seeded with HT115 bacteria and incubated at 15°C. After a recovery period of 72 h, the number of arrested animals at the L1/L2 stages and animals that developed beyond the L2 stage were counted. Survival percentages was calculated by dividing the number of arrested larvae by the total amount of animals.

## Statistics

Prism GraphPad was used to calculate statistical differences. For the survival experiments in Figure 3E and F a Log-rank (Mantel-Cox) test was used and for statistical significance between groups one-way ANOVA followed by post hoc analysis by Bonferroni’s test was used.

## Acknowledgements

We thank Dr. A. Theil for advice, Dr. G. Jansen for use of his injection microscope, Dr. Heng-Chi Lee and Dr. Andrew Fire for strains and plasmids and Dr. G. van Cappellen and the Erasmus MC Optical Imaging Center for microscope support. Some strains were provided by the Caenorhabditis Genetics Center (funded by NIH Office of Research Infrastructure Programs P40 OD010440) and the National Bioresource Project for the nematode. This work was supported by the Netherlands Organization for Scientific Research (711.018.007 and ALWOP.494), the Marie Curie Initial Training Network ‘‘aDDRess’’ funded by the European Commission 7th Framework Programme (316390), the European Research Council (advanced grant 340988-ERC-ID), and the gravitation program CancerGenomiCs.nl from the Netherlands Organization for Scientific Research. Oncode Institute is partly financed by the Dutch Cancer Society.

## Author contribution

KLT, MvdW, CD, MS and HL performed experiments and analyzed data. KLT, MvdW, WV and HL designed experiments. MvdW, WV and HL wrote the manuscript. All authors reviewed the manuscript.

