## Supplemental Material for "*C. elegans* TFIIH subunit GTF-2H5/TTDA is a non-essential transcription factor indispensable for DNA repair"

### Expanded view figure 1

**A**

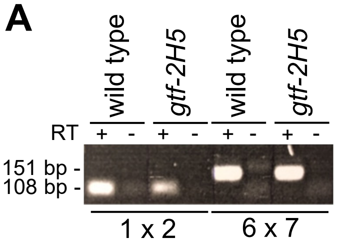

**B**

| primer pair | wild type | *gtf-2H5* |
|---|---|---|
| 1 x 2 | ++ | + |
| 1 x 3 | ++ | - |
| 1 x 4 | - | + |
| 1 x 5 | - | - |
| 2 x 6 | - | - |
| 6 x 7 | ++ | ++ |

**C**

```
                10        20        30        40        50        60
1    ATGGTGAACGTGAAGAAGGGAGTGTTGGTGACGAGTGATCCGGCTTTTCGTCAACTTCTG
1     M  V  N  V  K  K  G  V  L  V  T  S  D  P  A  F  R  Q  L  L

                70        80        90       100       110       120
61   ATCCATTTGGACGACTCCAGACAGCTCGGATCCAAGTTCATTGTCAGGGCACTTGTGCCA
21    I  H  L  D  D  S  R  Q  L  G  S  K  F  I  V  R  A  L  V  P

               130       140       150       160       170
121  CGTGGCGCAAGATGCAACGCATCGAGTTTCGCATTGGATTCCCGAATGTAG
41    R  G  A  R  C  N  A  S  S  F  A  L  D  S  R  M
```

**D**

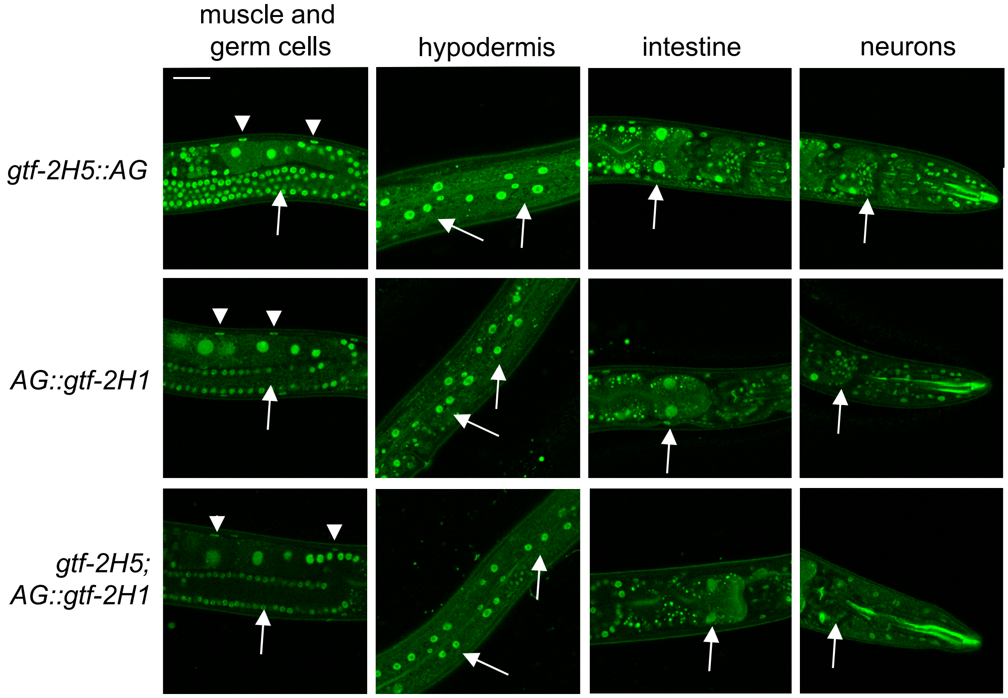

**Figure EV1 – Mutant *gtf-2H5* cDNA and expression of GTF-2H5 and GTF-2H1**

A. Independent RT-PCR from Fig 1A, confirming reduced levels of cDNA in *gtf-2H5* animals and showing that in these animals the mRNA of upstream DNA repair gene *helq-1* is normally produced, ruling out that any phenotypes observed in *gft-2H5* animals are due to an effect on HELQ-1.

B. Table to indicate which cDNA PCR products were detected with primer pairs shown in Fig 1 in wild type and *gtf-2H5* animals. '++' indicates that a strong and '+' that a weak PCR band was detected.

C. Shown is the sequence of the ORF found in the cDNA of the *tm6360* allele. Black nucleotide sequences are from the first exon of *gtf-2H5* while red colored sequences are reverse sequences derived from the B0353.1 gene. The putative translation is also indicated.

D. Representative confocal images showing fluorescence of GTF-2H5::AG (upper panels) and AG::GTF2H1 in wild type background (middle panels) and *gtf-2H5* background (lower panels). Tissues indicated are muscle (arrow heads) and germ cells, epidermal cells, intestinal cells and head neurons (arrows). Scale bar: 25 µm.

### Expanded view figure 2

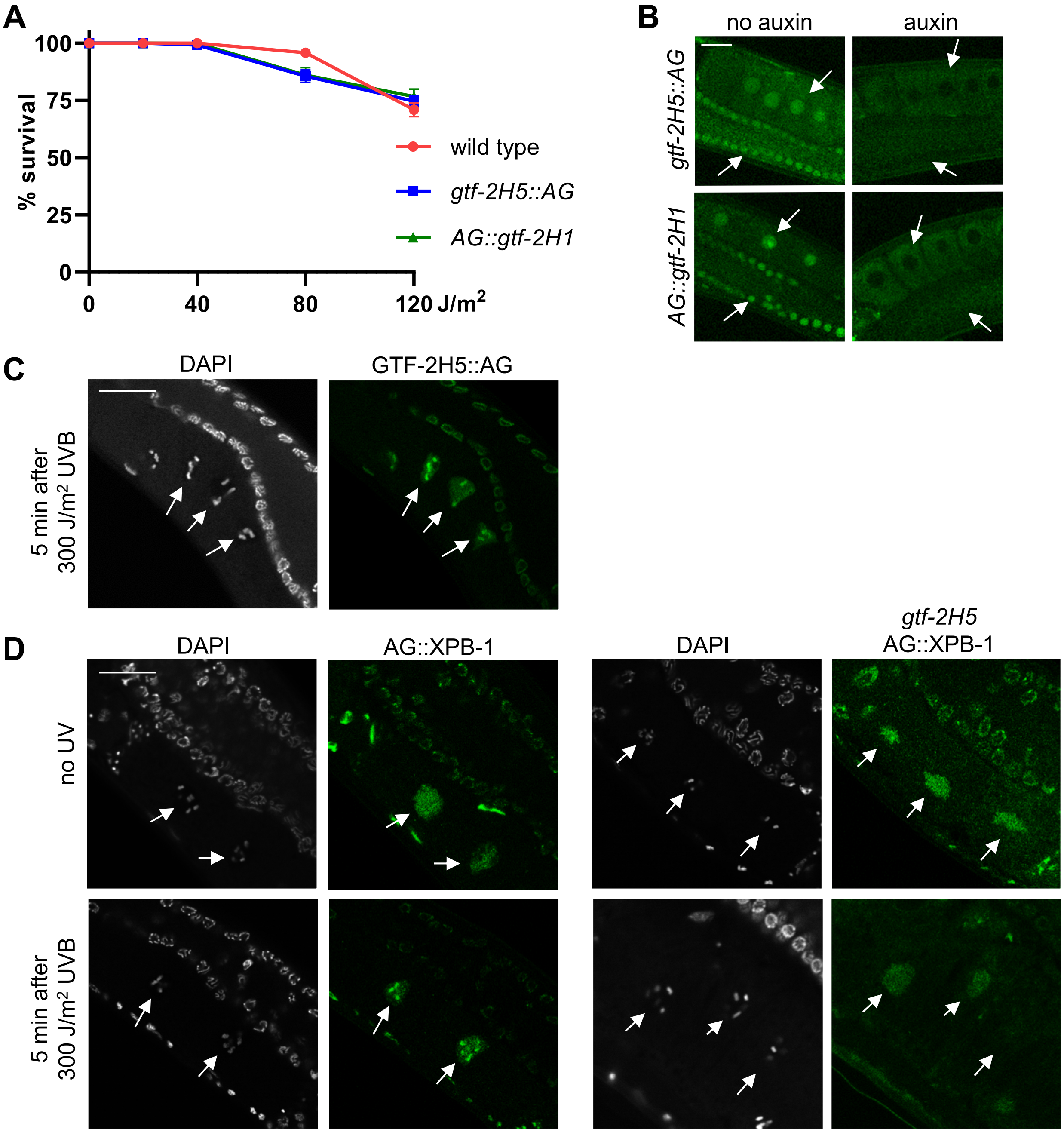

**Figure EV2 – *gtf-2H5::AG* and *AG::gtf-2H1* knock-in and TFIIH recruitment to DNA damage**

A. L1 larvae survival assay after UVB irradiation of wild type and *gtf-2H5::AG* and *AG::gft-2H1* knock-in animals. The percentages of animals that developed beyond the L2 stage (survival) are plotted against the applied UVB doses. Results are plotted as average with SEM (error bars) of ten experiments.

B. Representative confocal images showing efficient depletion of AID- and GFP-tagged GTF-2H1 and GTF-2H5 in animals expressing TIR1 under control of the *sun-1* promoter grown on 1 mM auxin for 24 h. Arrows indicate nuclei of germ cells. Scale bar: 20 µm.

C. Representative confocal images showing recruitment of GTF-2H5::AG to UVB-damaged chromosomes, visualized with DAPI (indicated in white), in oocytes of fixed wild type animals, 5 min after 300 J/m$^2$ UVB irradiation. Paired homologous chromosomes in oocytes are indicated with arrows. Scale bar: 20 µm.

D. Representative confocal images showing nuclear localization of AG::XPB-1 to UVB-damaged chromosomes, visualized with DAPI (indicated in white), in oocytes of fixed wild type (left) and *gtf-2hH5* (right) animals, without UV and 5 min after 300 J/m$^2$ UVB irradiation. Paired homologous chromosomes in oocytes are indicated with arrows. Scale bar: 20 µm.

#### Expanded view Table 1

| strain | genotype |
|---|---|
| CA1199 | *ieSi38[[P(sun-1)::TIR1::mRuby] IV* |
| CA1202 | *eSi57[P(eft-3)::TIR1::mRuby] II; ieSi58 [P(eft-3)::AID::GFP] IV* |
| GJ1566 | *xpa(ok698)I 6xoc* |
| HAL94 | *gtf-2H5(tm6360) III* |
| HAL203 | *gtf-2H5(tm6360) xpb-1(emc58[AID::GFP::xpb-1]) III* |
| HAL204 | *xpb-1(emc58[AID::GFP::xpb-1]) III* |
| HAL237 | *gtf-2H5(emc73[gtf-2H5::AID::GFP]) III; ieSi38[P(sun-1)::TIR::mRuby] IV* |
| HAL240 | *gtf-2H5(emc73[gtf-2H5::AID::GFP]) III* |
| HAL242 | *gtf-2H5(tm6360)III; gtf-2H1(emc202[AID::GFP::gtf-2H1]) IV* |
| HAL243 | *ieSi38[P(sun-1)::TIR1::mRuby] gtf-2H1(emc202[AID::GFP::gtf-2H1]) IV* |
| HAL254 | *ieSi57 [P(eft-3)::TIR1::mRuby] II; gtf-2H5(tm6360)III; ieSi58 [P(eft-3)::AID::GFP] IV* |
| HAL504 | *gtf-2H1(emc202[AID::GFP::gtf-2H1]) IV* |
| N2 | wildtype |

#### Expanded view Table 2

| | gene | sequence | sequence | primer Fig 1 |
|---|---|---|---|---|
| **RT-PCR** | *gtf-2H5* | GGTGAACGTGAAGAAGGGAG | | 1 |
| | *gtf-2H5* | CCTGACAATGAACTTGGATC | | 2 |
| | *gtf-2H5* | GTCAACAGCCTCCGGGATTC | | 3 |
| | *B0353.1* | GTTGCATCTTGCGCCACGTG | | 4 |
| | *B0353.1* | GAGCTCGCCTACTATCAATTG | | 5 |
| | *helq-1* | CAACGACTTTCTGAAGCTGCC | | 6 |
| | *helq-1* | CCCAAACGTCGAGTTCTTCC | | 7 |
| **RT-qPCR** | *cdc-42* | TCCACAGACCGACGTGTTTC | AGGCACCCATTTTTCTCGGA | |
| | *pmp-3* | GTTCCCGTGTTCATCACTCAT | ACACCGTCGAGAAGCTGTAGA | |
| | *gtf-2E1* | GGTGGATGAGATTCCGGAGG | GCTCGCATTATGTGGTAGACG | |
| | *gtf-2H5* | GGTGAACGTGAAGAAGGGAG | CCTGACAATGAACTTGGATC | |
